# Landscape and meteorological variables associated with *Aedes aegypti* and *Aedes albopictus* mosquito infestation in two southeastern USA coastal cities

**DOI:** 10.1101/2024.06.06.597792

**Authors:** Andre Luis Costa-da-Silva, Kyndall C Dye-Braumuller, Helen Urpi Wagner-Coello, Huixuan Li, Danielle Johnson-Carson, Sarah M Gunter, Melissa S Nolan, Matthew DeGennaro

## Abstract

*Aedes* transmitted arboviral human cases are increasing worldwide and spreading to new areas of the United States of America (USA). These diseases continue to re-emerge likely due to changes in vector ecology, urbanization, human migration, and larger range of climatic suitability. Recent shifts in landscape and weather variables are predicted to impact the habitat patterns of urban mosquitoes such as *Aedes aegypti* and *Aedes albopictus*. Miami (FL) is in the tropical zone and an established hotspot for arboviruses, while Charleston (SC) is in the humid subtropical zone and newly vulnerable. Although these coastal cities have distinct climates, both have hot summers. To understand mosquito infestation in both cities and potentiate our surveillance effort, we performed egg collections in the warmest season. We applied remote sensing with land-use cover and weather variation to identify mosquito infestation patterns. Our study found predominant occurrence of *Ae. aegypti* and, to a lesser extent, *Ae. albopictus* in both cities. We detected statistically significant positive and negative associations between entomological indicators and most weather variables in combined data from both cities. For all entomological indices, weekly wind speed and relative humidity were significantly positively associated, while precipitation and maximum temperature were significantly negatively associated. *Aedes* egg abundance was significantly positively associated with open land in Charleston but was negatively associated with vegetation cover in combined data. There is a clear need for further observational studies to determine the impact of climate change on *Ae. aegypti* and *Ae. albopictus* infestation in the Southeastern region of the USA.

## INTRODUCTION

Mosquito vectors are capable of transmitting pathogens that cause >700,000 deaths annually (WHO 2020). As mosquito-borne diseases continue to spread and reemerge, there is an increasing range of climate suitability, directly predicted to raise the number of people at risk to 4.7 billion by 2070 (Colón-González et al. 2021). Climate change can affect the transmission dynamics of vector-borne diseases by modifying the landscape and meteorological factors needed for mosquito survival and habitat suitability (Brugueras et al. 2020). The geographical expansion of mosquito-borne diseases driven by climate change is likely to intensify disease outbreaks globally, and local studies are urgently needed to understand the link between climate and arbovirus transmission (Robert, Stewart-Ibarra, and Estallo 2020).

Disproportionately affected by climate change, urban areas are at a greater risk of mosquito-borne diseases (Ligsay, Telle, and Paul 2021). Urban landscapes are more susceptible to ‘heat islands’—small geographic areas with up to ^+/-^7^○^F temperature variance—due to anthropogenic structures absorbing and re-emitting more heat (Huang et al. 2008; Mohajerani, Bakaric, and Jeffrey-Bailey 2017; Hibbard et al. 2017; EPA 2024). It has been shown that heat islands in urban areas have a higher arboviral disease incidence (Araujo et al. 2015). Further, over 90% of all urban areas are located in coastal cities which are more susceptible to flooding and extreme weather (Ligsay, Telle, and Paul 2021). These events can significantly impact urban availability of aquatic habitats for mosquito oviposition and larval development (Ligsay, Telle, and Paul 2021). Hurricanes in urban coastal cities are particularly concerning, as they can exacerbate and accelerate mosquito abundance and disease transmission potential (Lehman et al. 2007; Nasci and Moore 1998; AMCA 2022; Connelly and Borchert 2020).

While Miami, FL and Charleston, SC are two southeastern, coastal cities in the USA that experience similar hot summer climates (McGahan 1909; Merschman et al. 2020), they represent different seasonal weather patterns and land use/classifications. Specifically, Miami has a tropical wet/dry season with new construction and Charleston has humid subtropical seasons with preserved historic construction (Alamdari et al. 2018; Shulman et al. 2022; Wells and Baldwin 2012; MDC, n.d.; Charleston, n.d.). Miami-Dade County is an established hotspot for transmission of vector-borne diseases (Coatsworth et al. 2022; McAllister et al. 2020). It is the most populous county in Florida, where rapid urbanization processes and acute population growth changes have favored urban mosquito abundance, richness, and community composition (Wilke et al. 2019). Moreover, tourism in Miami-Dade provides a constant connection to arbovirus influx from endemic regions, especially Central and South America (Alonso 2007). Conversely, Charleston County has historically been a hotspot for arbovirus outbreaks (Beaumier, Garcia, and Murray 2014; Duffy 1951), yet poses arboviral outbreak vulnerability due to similar environmental conditions and a vibrant Caribbean island cruise industry port (Magill 2014; McLeod 2013; SC Ports Authority 2015; Marsh 2012). Therefore, to better understand the dichotomy of arboviral disease transmission risk between an emerging-endemic American coastal city and an epidemic at-risk coastal city, we performed mosquito egg collections in urban areas of Miami, FL and Charleston, SC. Our primary goal was to understand *Aedes aegypti* and *Aedes albopictus* infestation and corresponding climate and environmental risk factors for *Aedes* mosquito abundance in these two communities.

## MATERIALS AND METHODS

### Study areas

Ovitrap egg collections were performed weekly from June - August 2019 in Miami-Dade County (FL) and Charleston County (SC). Sampling was predominantly in urban, high-developed areas of both counties. Miami-Dade County is a large, rapidly growing metropolis, home to 2.7 million residents and new construction, especially within the urban center (US Census 2022). In contrast, Charleston County is home to a moderate-size city of 419,000 residents with centuries-old buildings and considerable historical preservation ordinances limiting new construction in the city (US Census 2022).

### *Aedes* Mosquito Egg Collection

Ovitrap styles and deployment methods were similar for both locations. Ovitraps consisted of a black plastic cup [112 mm height x 92 mm top diameter x 70 mm base diameter (450 mL capacity)]. A drainage hole of 13 mm diameter on the lateral side of the cup (at 65 mm height) was drilled to avoid overflow during rain events. Ovitraps were labeled with participating organizations’ contact information to prevent misplacement. The ovitrap was lined with a strip of 40 mm length x 75 mm width of heavy weight seed germination paper (#SD7615L, Anchor Paper Co.) and filled with 6.76 fl oz (200 mL) of tap water. After the deployment period, the paper sample recovered from the ovitrap was placed inside a labeled Whirl-Pak® bag (Nasco Sampling/Whirl-Pak, Madison, WI, USA), and trap water was discarded. The samples were transported to the laboratory and germination papers were allowed to air dry for 24 hours. The dried papers were screened under a dissection scope and mosquito eggs were counted by the collecting university partner. All eggs were considered regardless of the condition (intact, hatched, unhatched, dried, or broken).

The positive samples were hatched by transferring the seed paper into a plastic rearing pan (5.3 L polycarbonate food pan, Carlisle, Oklahoma, USA) containing 2L of DI water. The hatched larvae were fed with TetraMin tropical tablets (Tetra, Melle, Germany) until eclosing in adults. The samples from Charleston had a low hatchability. We speculate that this may be due to the shipment of the samples from Charleston to Miami, the site where the samples were processed. The pupae were transferred to cages and eclosed adult mosquitoes were cold-anesthetized for species identification under a stereoscopic microscope (Olympus SZ51, Olympus Corporation, Tokyo, Japan). The species identification was performed following the identification keys previously described (Consoli and Oliveira 1994).

### Miami-Dade County Ovicup Trapping Locations and Sampling

Locations for Miami urban areas were based on the residential areas available from volunteers of the Florida *Ae. aegypti* Genome Group (FLAGG) internship coordinated by the DeGennaro Laboratory of Tropical Genetics at Florida International University. This internship was open to all university students in summer 2019. Volunteer students received the collection kits to set up the ovicups at their residences over the summer. Kits included two ovitraps, strips of seed germination paper, and a Whirl-Pak® bag for sample collection. Ovitraps lined with a strip of seed germination paper and filled with tap water were deployed weekly in 43 residential sites in and around Miami-Dade County from June to August 2019. Ovitraps were placed in shaded areas of residential yards on Thursday mornings and collected on Sunday afternoons (Fonseca et al. 2013). A 3-day exposure time was determined to mitigate larval development given Miami’s warmer climate.

### Charleston County Ovicup Trapping Locations and Sampling

Mosquito site selection in South Carolina occurred through recommendations by the local vector control agency (Charleston County Mosquito Control) and manual field site determination across the Charleston peninsula. During May 2019, multiple churches with cemeteries, local parks, elementary schools, independently owned businesses, and the Citadel Military College of South Carolina were contacted for permission to place ovitraps in shaded areas for the June - August 2019 sampling duration. To represent diverse land-use classifications, 35 sites were ultimately selected. As in Miami, the same ovitraps, germination paper, and tap water were used in Charleston. Weekly, ovitraps were placed in shaded areas (Fonseca et al. 2013) under bushes or trees (specifically oaks), or adjacent to tires when available. Ovitraps were deployed on Mondays in the morning, and germination papers were collected on Fridays.

### Climatic and Environmental Variables

Weekly climate variables, including temperature, relative humidity, wind speed, and precipitation, were recorded during the study period from the ovitraps’ nearest weather stations [National Climatic Data Center’s Climate Data Online tool (National Centers for Environmental Information, n.d.)]. Sentinel-2 satellite images were obtained from the European Space Agency and imported into a U-Net convolutional neural network deep-learning algorithm to land-use land-cover classification variables. Additional indices for water content (NDWI) and vegetation (NDVI) were derived from the same images.

### Geographical Analyses

Latitude and longitude coordinates were mapped for each unique ovitrap site, and a 300m^2^ buffer was created around each coordinate to clip and extract relevant land-use land-cover data for each location. Land use and land cover data was extracted from the National Oceanic and Atmosphere Administration C-Cap 10m as percent prevalence within a buffer or as the median index value within the buffer. The trap location diameter was selected to accommodate for urban mosquito species’ flight ranges (Nagpal et al. 2016). The maps for both Charleston and Miami-Dade County collections categorized by OI were produced using ArcGIS Pro (ESRI, Redlands, CA).

### Statistical Analyses

Ovitrap indices (Egg Density Index from positive traps - EDI^P^, Egg Density Index from all traps deployed - EDI^T^, and Oviposition index - OI) were calculated by site and epidemiological week. EDI^P^ estimates the intensity of infestation levels (total number eggs recovered/total positive ovicups), EDI^T^ indicates the distribution of the infestation (total number eggs recovered/total ovicups deployed), and OI estimates the rate of positive traps (number of positive ovicups/total ovicups deployed). Descriptive statistics were calculated for all land-use land-classification variables for both cities. Before statistical analyses, approximate normality was achieved by a square root transformation of each egg index. Univariate regression models were performed for each variable utilizing egg abundance and each egg index separately; simple linear regression was used for each ovitrap index, while a negative binomial regression was used for the egg abundance outcome variable only to address egg count data with overdispersion. To account for multiple comparisons, all raw p-values were corrected with the Bonferroni method. All statistics were performed using SAS 9.3 (SAS Institute, Cary, NC, USA).

## RESULTS

Ovicups were deployed in Charleston (35 sites) and in Miami-Dade County (43 sites) over 10 epidemiological weeks during the 2019 summer season. The oviposition index (OI) for each site was obtained and mapped for both cities (Figure 1). We observed varied OI ranges among the sites for each city, but only Miami showed negative sites for eggs over the course of the study (8 sites, OI=0) (Figure 1). Out of 35 positive sites in Miami, samples from 31 sites hatched. *Ae. aegypti* was found in 30 sites and *Ae. albopictus* was identified in only 1 site Charleston had 35 positive sites, but only samples from 4 sites hatched. *Ae. aegypti* was identified in 3 sites and 1 site showed *Ae. albopictus*.

**Figure 1.**
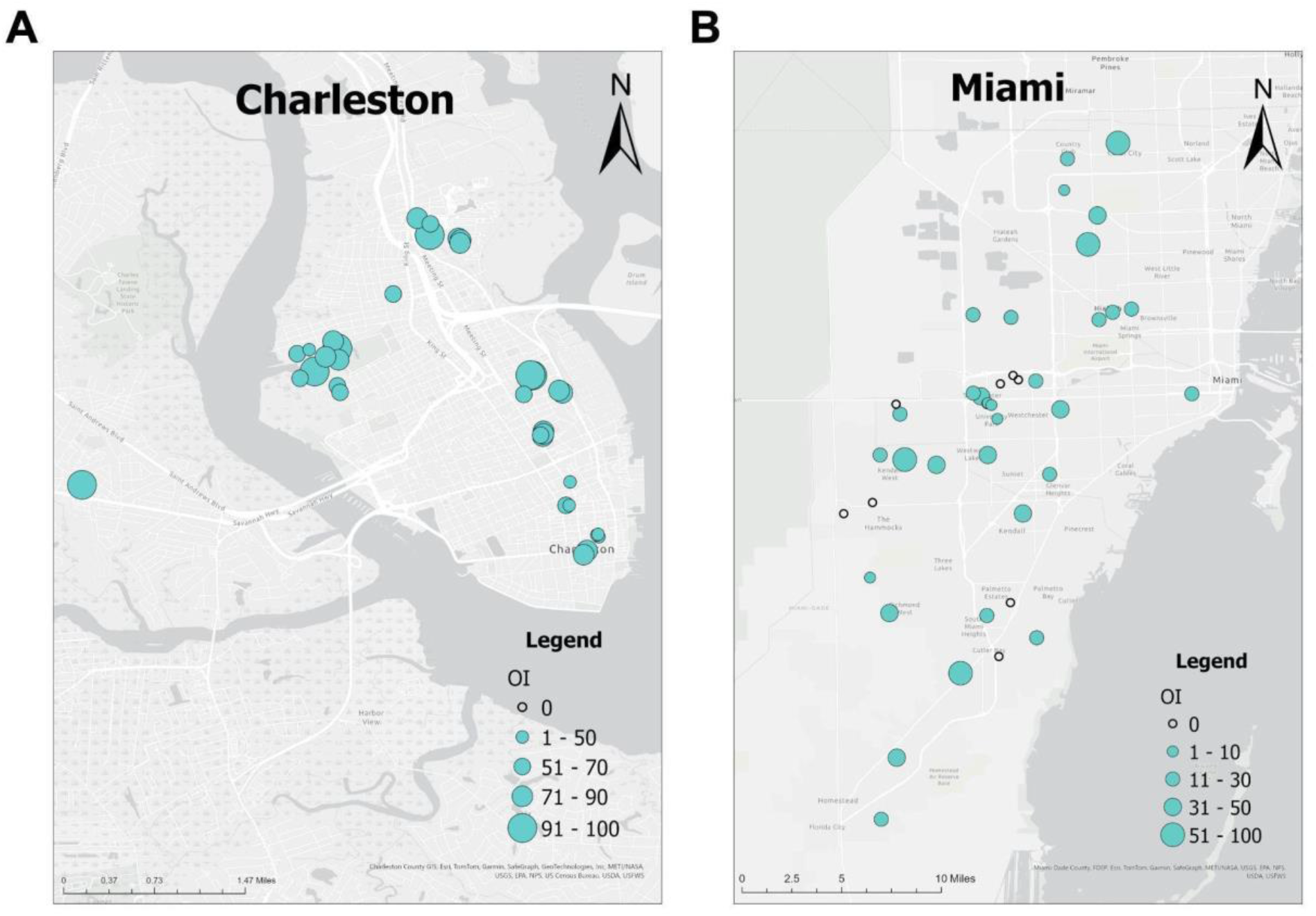
Oviposition index of the collection sites in Charleston, SC (A) and Miami, FL (B) over the 2019 summer season. Sites with OI of 0 are indicated with a black outlined dot. Sites with OI higher than 0 are indicated by green dots. The OI ranges are in accordance with the legends embedded in the figure.

Both cities had similar weather conditions throughout summer season 2019. However, Charleston was cooler (avg. range: 87.1-76.8°F vs. 92.0-77.1°F), had a higher relative humidity (74.8% vs. 70.4%), higher wind speeds (6.6 mph vs. 5.8 mph), and less precipitation (0.1 in. vs. 0.4 in.) compared to Miami (Figure 2).

**Figure 2.**
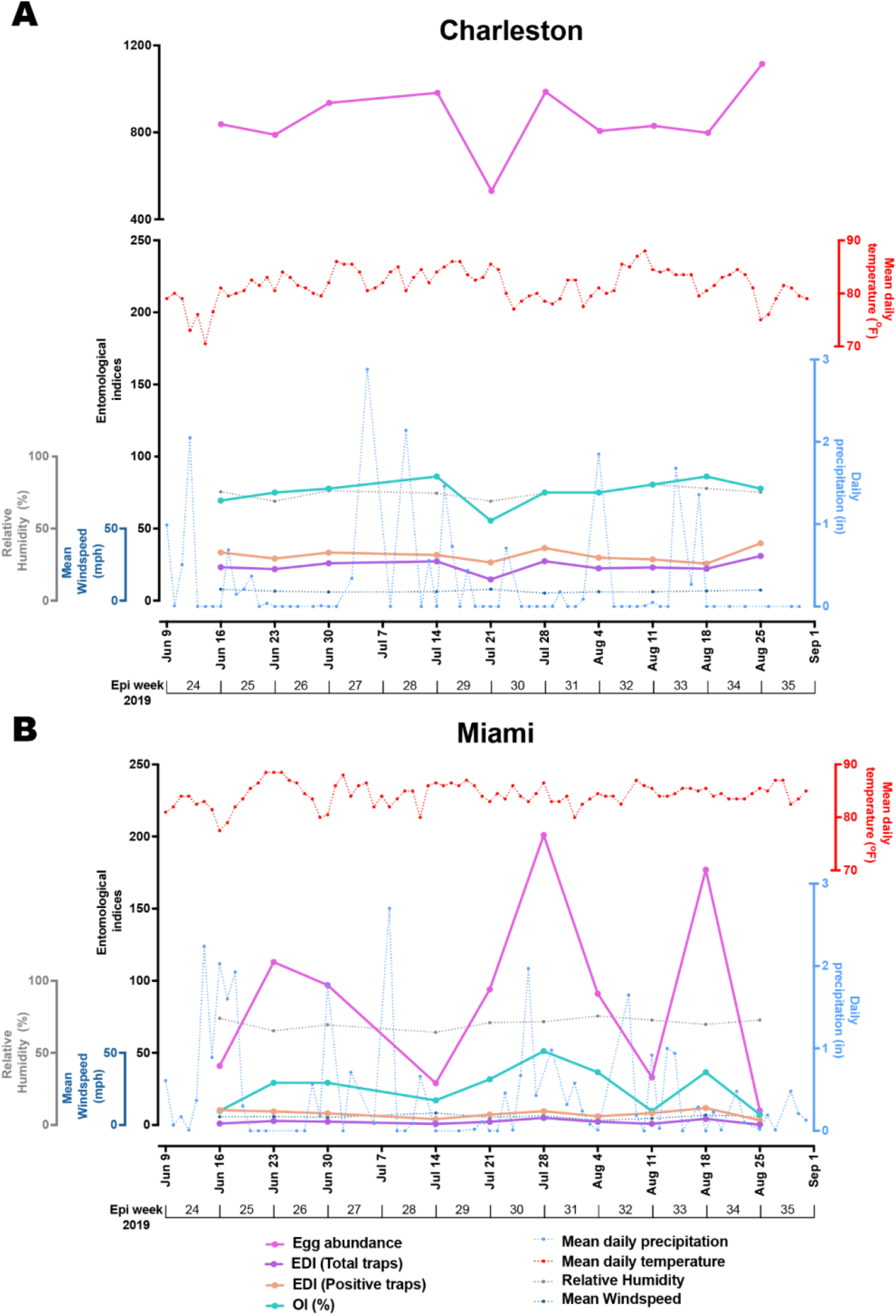
Timeline comparison of meteorological variables, egg abundance and entomological indices of mosquito eggs in (A) Charleston (SC) and (B) Miami (FL) during summer 2019. Weekly egg abundance (pink line), egg density index from the total traps (EDI^T^, purple line), egg density index from the positive traps (EDI^P^, orange line) and oviposition positivity index (OI, green line) compared to weekly mean precipitation (light blue-dotted line), temperature (red-dotted line), relative humidity (gray-dotted line), and mean wind speed (dark blue-dotted line) from June to September, 2019.

Egg abundance and entomological indices for the ovitrap collections varied between the two cities (Table 1 and Figure 2). Charleston, SC yielded a higher absolute number of total eggs and higher entomologic indices (EDI^P^, EDI^T^, and OI) than Miami, FL. Both cities’ 300m^2^ buffers around ovitraps had similar percentages of the different land classifications, with the highest proportion consisting of medium development (35.3% for Charleston and 37.3% for Miami). Additional common land classifications for Charleston were high development (30.1%), low development (22.8%), open land (9.0%), emerging herbaceous wetlands (2.1%), and water (0.2%). Conversely, Miami had less high development (11.4%) and greater low development (34.0%) plots around mosquito traps. Open land (10.5%), emerging herbaceous wetlands (0.9%) and standing water (1.8%) were similar in Miami. Low prevalent land classifications (<0.1%) were collapsed into a single “vegetation cover” variable to represent a collective ‘other’ grouping: including, bare land, deciduous forest, evergreen forest, shrub scrub, hay pasture, cultivated crops, and woody wetlands.

**Table 1.**
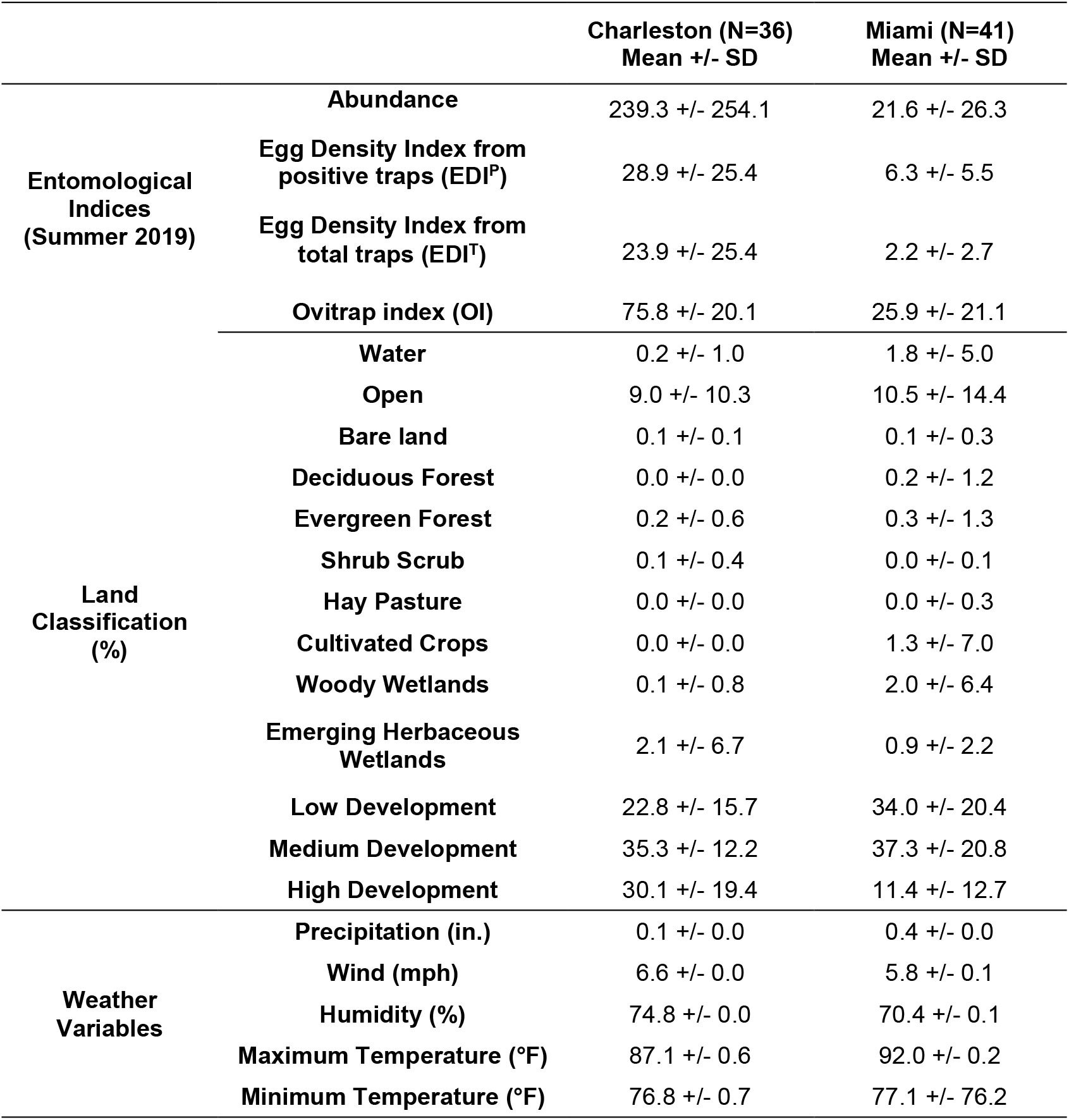
Descriptive Statistics on Environmental and Land Classification Variables by City.

Table 2 presents the simple linear regression univariate models (for all egg indices) and the negative binomial regression univariate models (for egg abundance only) results to identify environmental and land correlations within each city and combined. Only egg abundance was associated with a land classification variable in the negative binomial regression. Open land was positively associated with *Aedes* egg abundance in Charleston only. No weather variables were significant, and no other egg index was associated with any variables. Conversely, Miami demonstrated no *Aedes* egg count associations with neither weather variables nor land classification types.

**Table 2.**
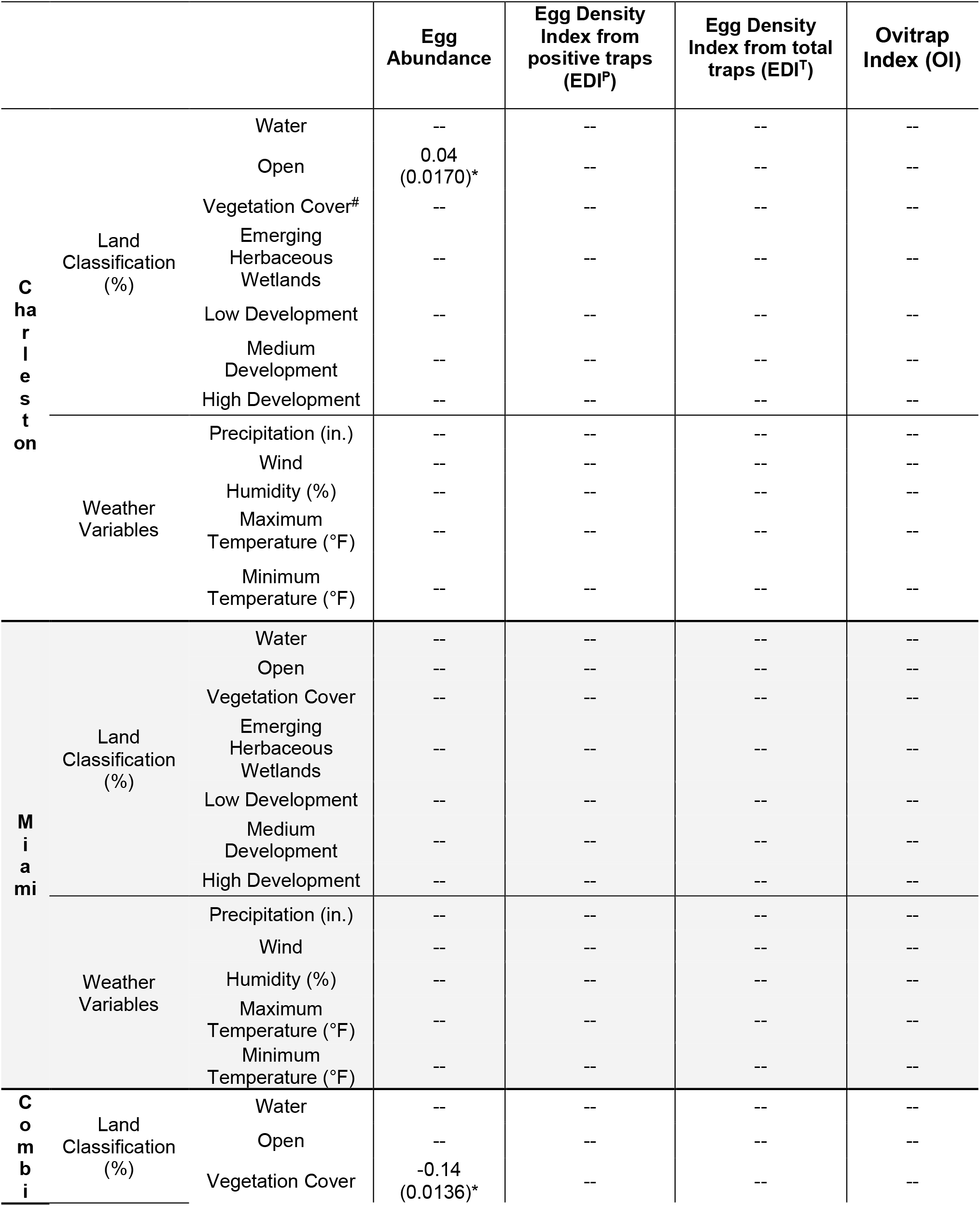
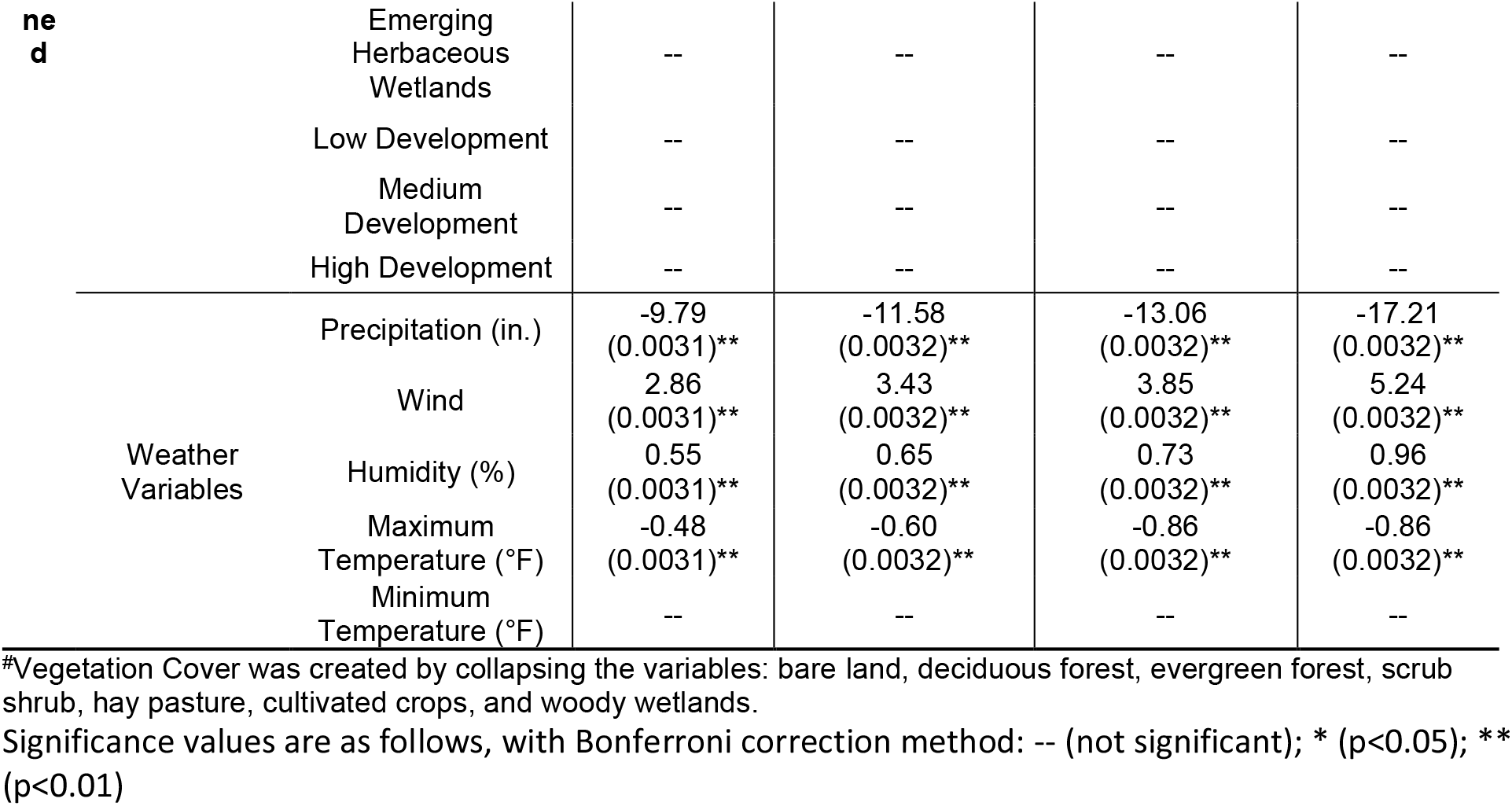
Univariate Regression Models with Different Egg Density Measurements.

Combined datasets (Charleston and Miami) yielded the highest number of statistically significant variables, particularly among weather factors. In combined models, presence of vegetation cover (heterogeneous mix of rare land cover classification types) was significantly negatively associated with egg abundance. Weather variables for the combined model were statistically significant across all four container-breeding mosquito egg measurements: precipitation, wind, humidity, and maximum temperature. Notably, all egg outcome variables were reduced with increases in precipitation and ambient temperature. Conversely, all egg outcome variables were increased with increases in wind speed and by relative humidity.

## DISCUSSION

This unique study evaluated the distribution and environmental-climatic variables associated with mosquito egg abundance and infestation indices in two ecologically different, coastal Southeastern USA cities over the same period. *Ae. aegypti* and *Ae. albopictus* were the only two species detected regardless of the city. We chose to deploy ovicups due to the high efficiency of this simple trapping device in vector surveillance of container-breeding mosquitoes (Djiappi-Tchamen et al. 2022; Martini et al. 2017; Mardiana et al. 2018). Differently from autocidal gravid ovitrap (AGO trap) (Barrera et al. 2014), ovicups cannot be exposed for longer periods to prevent the creation of productive breeding sites. Even though a short period of ovicup exposure was defined in our study, positive samples were frequent during the collection period. In addition, it has been shown that the ovitrap positivity index can be more sensitive than the survey of immature stages (Wijegunawardana et al. 2019; Djiappi-Tchamen et al. 2021; Rawlins et al. 1998; Braga et al. 2000).

The use of the ovicup method and estimation of egg abundance and infestation indices allowed us to detect significant associations with weather variables when data from both cities was combined, and a positive association between infestation and landscape for one of the cities. Charleston, SC was founded in the late 1600s and historical buildings predominate today; whereas Miami, FL was largely established in the mid-1900s with modern construction commonplace. Interestingly, separately, Charleston was the only city with an associated land classification variable with egg abundance—a surprising finding as Miami is a well-known location of human container-breeding (*Aedes* spp.) mosquito transmitted pathogens. While distinctions in container-breeding mosquito abundance factors were identified in each ecological area, some important commonalities existed—both aspects presenting implications for generalizable and tailored vector control interventions (Lindsey et al. 2012; CDC, NCEZID, DVBD 2023).

Both cities are tourist destinations with notable recent expansion that has created rapid environmental changes (Rifat and Liu 2022; Transue et al. 2023; Rifat and Liu 2019). — Specifically, Charleston’s population grew 17.6% from 2010-2019, averaging an approximate increase of 1.95% annually, while Miami’s population grew 23.8% from 2000 to 2018, averaging an approximate increase of 1.32% annually (Transue et al. 2023; Rifat and Liu 2019). Interestingly, land development (increased population density and city infrastructure growth) was not associated with increased egg abundance in both cities. Traditionally, the abundance of urban mosquitoes, such as *Ae. aegypti* and *Ae. albopictus* species, increases in areas of high land development with warm tropical climates (Li et al. 2014; Wilke et al. 2020). It is possible that urban microclimates created by focal urban heat islands were inhibitory to local *Aedes* mosquito populations, as summer temperatures in these cities can typically exceed the preferred ranges for *Ae. aegypti* and *Ae. albopictus* (Reinhold, Lazzari, and Lahondère 2018). Therefore, the positive open land association in Charleston might indicate a byproduct of container *Aedes* populations migrating away from urban heat islands. However, the discrepancy observed with the lack of association between this landscape and egg abundance in Miami, suggests that this effect may not be a common occurrence when drastic landscape differences are developing in the same landscape classification (Divya et al. 2021; Wells and Baldwin 2012).

Urbanization, when rapid, poorly planned or managed, can increase the presence of container-*Aedes* mosquito larval habitats, such as debris containers and puddles in open land areas (Souza et al. 2023). Unlike developed urban areas with robust infrastructure and drainage systems, vacant and abandoned open land may receive limited human intervention. This can lead to any present containers, debris, or natural depressions that can hold undisturbed water (Haase, Haase, and Rink 2014). Open lands can function as peridomestic environments, subject to the influence of urban activities that can create artificial breeding sites for peridomestic species (Baak-Baak et al. 2014; Little, Bajwa, and Shaman 2017; Macêdo et al. 2021; Goldstein, Jensen, and Reiskin 2001). To understand the association between open land classification and mosquito presence and abundance, land management practices and urban development in open land areas should be further investigated.

Vegetation was additionally statistically negatively associated with moderately lower container-breeding mosquito egg measures in our combined dataset. Urban environments include a mix of vegetation types and range from highly maintained parks and gardens to overgrown vacant lots or poorly managed green spaces (Jansson and Randrup 2020). Well-maintained parks with proper drainage may minimize areas of standing water and support less mosquito breeding. Meanwhile, neglected or poorly managed green spaces might provide ideal mosquito breeding conditions through stagnant water and debris (Obame-Nkoghe et al. 2023). Although peridomestic species, such as *Ae. albopictus*, can take advantage of some levels of vegetation immersed in urban areas (O’Meara et al. 1993), a higher level of vegetation coverage limits the occurrence of mosquito urban species and their oviposition activity in ovicups (artificial breeding sites) (Benitez et al. 2020).

Lastly, the combined city model revealed a strong climate influence on relative mosquito abundance. Specifically, precipitation and temperature were negatively associated with container-breeding mosquito egg measures, whereas wind and humidity were positively associated. As previously mentioned, these cities routinely experience sustained temperatures above *Aedes* preferred temperature range, which may explain the low egg yield/capture during higher temperature times. The negative association with rainfall has been observed previously and it might suggest competition among new rain-produced breeding sites (Resende et al. 2013). Further, heavy rainfall can flush out mosquito larvae from breeding sites, temporarily reducing local populations (Paaijmans et al. 2007). Conversely, higher humidity is a known physiological benefit for *Aedes* mosquitoes to prevent desiccation and both cities’ respective humidity levels averaged within the optimum activity ranges (Manzano-Alvarez et al. 2023; Wei et al. 2023; Cai et al. 2023).

Wind speed and mosquito prevalence are understudied correlations, and our study is one of the first to describe a statistically significant positive correlation between increased wind speeds and mosquito egg abundance. For adult mosquito populations, wind speeds and direction can play a facilitative role in mosquito movement and foraging activities by aiding their dispersal and navigation (Service, M. W 1997; Ferreira et al. 2006; Hribar, DeMay, and Lund 2010). The wind pushes the mosquitoes, allowing them to discover new breeding sites and hosts for blood meals over a greater reach radius, potentially leading to increased mosquito abundance (Hribar, DeMay, and Lund 2010). Alternatively, it is possible that higher wind speeds can prevent gravid female mosquitoes from flying away from the ovicup or seeking more sites when skip-ovipositing (Day 2016; Atieli et al. 2023). Limited scientific literature describes lower adult mosquito flight efficiency, which contributes to the hypothesis that gravid females might be more likely to lower their standards for egg laying selection sites in windy conditions (Day 2016; Atieli et al. 2023).

The southeastern USA is undergoing concomitant population expansion, urbanization and climate change induced environmental effects that support rapid container-*Aedes* spp mosquito population migration and national range expansion over the next few decades, as noted by a myriad of scientific predictions (Khan et al. 2020; Kache et al. 2020; Ryan et al. 2019; Hall et al. 2022; Gloria-Soria et al. 2020). This study performed a comparative analysis of container-breeding mosquito egg measures and ecology between two distinct southeastern USA coastal cities: Charleston, SC, and Miami, FL. Surprisingly, Charleston produced one associated land classification, whereas Miami had no detected associations with egg abundance or indices, despite Miami being a historical hot-spot for *Ae. aegypti* and *Ae. albopictus*. Land use and land cover categories had limited associations with mosquito abundance: namely areas with more open land yielded greater *Aedes* egg measures, and land use with mixed vegetation cover had a decreased association with container breeding mosquito eggs. Conversely, four climate variables had strong statistical correlations, suggesting weather might play a greater role in *Aedes* gravid female egg laying placement than gross environmental characteristics. These findings have both general and city-specific vector control intervention implications and warrant further study as ecologies change in the proceeding decades.

## Acknowledgments

The authors would like to thank the Charleston County Mosquito Control Program for helping select sites for *Aedes* mosquito breeding locations and allowing the use of their facilities throughout the summer, the work of McKenzi Norris, an undergraduate student at the time, for his help in deploying ovicups in Charleston throughout the duration of the project, and all cohort interns and volunteers that took part in the FIU FLAGG Internship in Summer 2019.

